# Clade C MERS-CoV camel strains vary in protease utilization during viral entry

**DOI:** 10.1101/2025.11.15.688481

**Authors:** Helena Winstone, Helen Stillwell, Tiffany Tan, Noam A. Cohen, Ranawaka A.P.M. Perera, Susan R. Weiss

## Abstract

MERS-CoV is a lethal pathogen with pandemic potential. Clade A and B MERS-CoV viruses have caused outbreaks in the Middle East since 2012 when they initially spilled over from camels to humans. Clade C viruses, however, are only found in camels across Africa and the spillover potential of these viruses seems to be lower than for clade A/B strains but remains to be fully understood. Here, we report that clade C spikes are less well-cleaved at the S1/S2 boundary than clade A or B viral spikes and that most clade C spikes induce reduced syncytium formation. Additionally, we demonstrate that several East African clade C strains are less able to utilize the TMPRSS2-mediated pathway for viral entry in both cell lines and primary nasal epithelial cultures. We map the molecular basis of this reduced TMPRSS2 usage to the N-terminal domain (NTD) and subdomain 2 (SD2) of East African clade C MERS-CoV. We suggest that reduced usage of the TMPRSS2-mediated entry pathway may underlie the reduced replication of East African clade C strains in humans, while the reduced replication of West African strains remains to be further investigated. Overall, we suggest that altered protease usage may contribute to differential tropism of East African clade C strains and indicate geographically distinct selection pressures on spike between MERS-CoV strains circulating in camels.

**Significance Statement:** Clade A/B MERS-CoV outbreaks have caused 957 deaths since the first spillover in 2012; meanwhile, Clade C MERS-CoV strains have been found in camels across Africa but have not been reported to cause outbreaks. Investigating why these viruses do not successfully transmit to humans will be key to understanding the pandemic potential of the African MERS-CoV camel reservoir. Our study indicates that clade C viruses exhibit less spike cleavage and that East African clade C isolates are less able to utilize the TMPRSS2-mediated pathway during viral entry of both human cell lines and primary nasal cells. Differences in viral entry pathways could alter cellular and organ tropism and inform our understanding of the pandemic potential of these viruses.

## Introduction

Middle East Respiratory Syndrome Coronavirus (MERS-CoV) is a betacoronavirus in the merbecovirus subgenus which has caused outbreaks in the Middle East since 2012. There have been over 2,627 cases as of April 2025 with a lethality rate of 35% [1]. Like SARS-CoV and SARS-CoV-2, MERS-CoV ancestral strains originated in bats; however, most human cases are attributed to infections acquired from interactions with the amplifying reservoir host, camels [2]. There are three clades of MERS-CoV which are geographically distinct: A, B, and C [3]. Clade A includes the prototypic Erasmus Medical Center (EMC) strain which caused the 2012 outbreak, however these viruses have since been outcompeted by clade B viruses [4]. Clade B viruses circulate in camels across the Middle East and cause frequent spillover events, with hundreds of cases reported in Saudi Arabia between 2013 and 2020 [5]. Clade C viruses, however, are found only in camels across Africa and there is currently no evidence of outbreaks attributed to these viruses. These viruses are phylogenetically distinct from clade A/B strains, with amino acid substitutions or deletions present in both the viral entry protein and key accessory genes previously shown to antagonize innate immunity [3].

MERS-CoV entry is mediated by the viral entry protein, spike, binding to the cellular receptor DPP4 [6]. MERS-CoV spike is a trimer of heterodimers which are composed of two subunits, S1 and S2. S1 contains the N-terminal domain (NTD), which has been shown to bind sialic acids and facilitate attachment [7, 8], and the receptor-binding domain (RBD), which binds to DPP4. The S2 subunit contains the fusion peptide which promotes fusion of the cellular and viral membranes following receptor engagement. Like several other coronaviruses, MERS-CoV spike requires activation in order to bind DPP4 in the form of proteolytic cleavage. Spike contains two cleavage sites, one at the S1/S2 boundary, and one at the S2’ site in the S2 subunit. The first cleavage event, at the S1/S2 site results in conformational changes that allow the RBD to engage DPP4. The second cleavage event at the S2’ site is required for further conformational changes which ultimately release the fusion peptide and initiate fusion of the viral and host membranes [9].

During viral biogenesis, the S1/S2 site can be cleaved by furin as spike trafficks through the Golgi. If not sufficiently cleaved, and if endosomal proteases are available, endosomal proteases during infection of the target cell can also cleave the S1/S2 site. It has been reported that both furin and cathepsin L can cleave both the MERS-CoV S1/S2 site and subsequently the S2’ site, allowing the virus to enter the cell via late endosomes. If sufficient S1/S2 cleavage has occurred during spike biogenesis, and TMPRSS proteases such as TMPRSS2 are available on the target cell surface, TMPRSS proteases can mediate the S2’ cleavage at the cell surface and the virus can fuse at the cell membrane [9–11]. Thus, relative protease bioavailability across cell-types and prior S1/S2 cleavage can alter the cellular tropism of MERS-CoV [11]. In addition to altering susceptibility to MERS-CoV infection, protease bioavailability has been reported to influence the efficiency of viral entry. For SARS-CoV-2, it has been reported that the “early” pathway of viral entry mediated by TMPRSS2 is faster (10-15 minutes) than the “late” pathway mediated by endosomal proteases (45-60 minutes) [12].

While the viral entry of clade A and clade B strains of MERS-CoV has been well characterized, to date the impact of amino acid substitutions in the spike gene of clade C strains on viral entry has not yet been fully examined. It has been confirmed that several clade C spike proteins do not differ from wild-type (WT) MERS in their ability to enter 293T cells overexpressing the receptor DPP4 as pseudotyped vectors [13]. However, differences in protease usage and entry into more physiologically relevant cell types have not yet been investigated. While none of the amino acid (aa) substitutions in clade C spikes are within DPP4-contacting residues or the S1/S2 cleavage site, previous studies of other coronaviruses’ spike have indicated that sequence changes outside the RBD can impact spike cleavage, protease usage, and cell-cell fusion [9, 14–17].

Here, we utilize a panel of six clade C MERS-CoV strains to demonstrate that clade C spikes are less cleaved at the S1/S2 cleavage site. Additionally, we report that East African clade C strains HKU270, CAC9690 and CAC10200 are less able to utilize the TMPRSS2-mediated pathway for viral entry. We identify that amino acid substitutions in the NTD, which are shared by all publicly accessible East African camel-derived MERS-CoV strains to date, confer reduced TMPRSS2 usage. Separately, the V677L substitution in the SD2 region of HKU270 spike also reduces TMPRSS2 usage. We suggest that these differences in viral entry pathway usage may contribute to differential tropism of these clade C viruses and indicate differential selection pressures on spike between MERS-CoV strains circulating in camels.

## Results

### Clade C viruses are attenuated for viral replication in A549^DPP4^ and Calu-3 cells relative to clade A/B viruses

To begin to determine factors in the differential spillover potential of Clade C isolates, we utilized six clade C isolates, three from West Africa (Nig1657, Mor213 and BF785) and three from East Africa (HKU270, CAC9690, CAC10200), along with WT EMC, a clade A strain, and AH13, a clade B strain. All the clade C viruses contain 6-8 amino acid substitutions in the spike gene (**Fig 1A**). None of these substitutions are in direct DPP4-contacting residues or the S1/S2 cleavage site; however, it has previously been reported that amino acid substitutions in the NTD, SD2, and S2 regions of spike can alter spike conformation, cleavage, and fusion [9, 14, 15, 17]. Mor213 does contain a mutation in the S2’ site, R884L. AH13, a clade B virus, also contains 5 amino acid substitutions relative to EMC; however, none of these are the same as those in the clade C spikes. We therefore hypothesized that clade C spike substitutions may alter clade C viral entry in the human airways. Viral stocks of these strains were generated and titered in Vero-CCL81 cells (**Fig 1B**). Interestingly, all of the clade C viruses exhibited a smaller plaque morphology in Vero-CCL81 cells than the wild-type EMC strain or the clade B strain AH13 (**Fig 1B).** This was particularly apparent for Nig1657, Mor213 and CAC9690. BF785 also exhibited a peculiar plaque phenotype, with large fuzzy plaques. Despite their different plaque morphology, most of the clade C viruses produced sufficient titer in Vero-CCL81 cells (**Fig 1C**). Mor213 and CAC9690, however, were reduced by 2-logs at peak replication in these cells.

**Figure 1.**
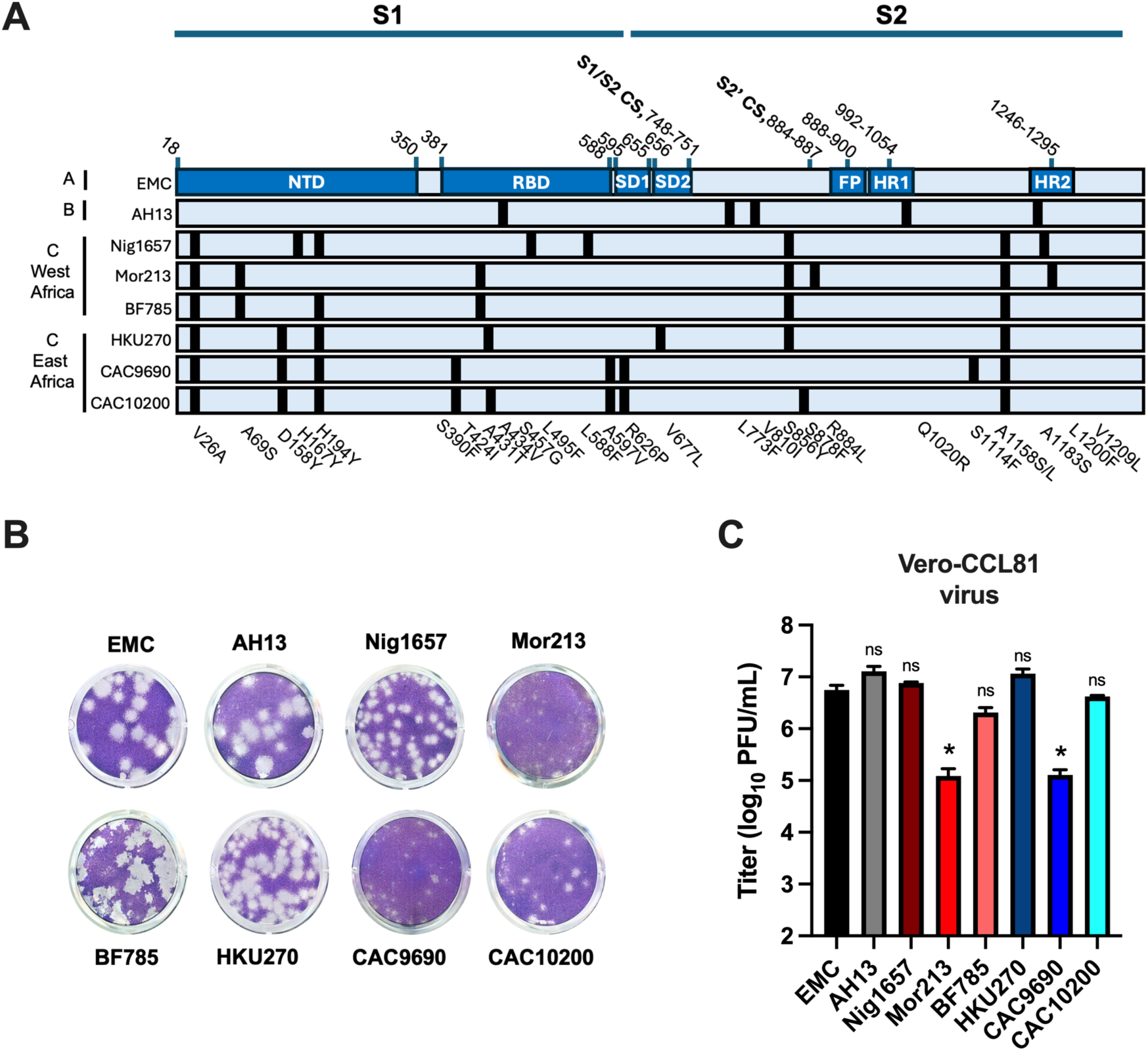
Clade C viruses contain amino acid substitutions in the spike gene. A) Amino acid changes found in clade B strain AH13 and six clade C viruses are shown. NTD = N-terminal domain, RBD = receptor binding domain, SD1 = subdomain 1, SD2 = subdomain 2, FP = fusion peptide, HR1 = heptad repeat 1, HR2 = heptad repeat 2. **B)** Representative images of plaque morphology of clade A, B and C strains in Vero-CCL81 cells. **C)** Vero-CCL81 cells were infected with viruses for 24h at MOI 0.1 and infectivity measured by plaque assay. N=3, *=P<0.05 tested by one-way ANOVA showing statistical significance against WT EMC.

Next, we examined the replication of this panel of viruses in respiratory cell lines.

A549^DPP4^ and Calu-3 cells were infected for 16, 24, or 48h with EMC (clade A), AH13 (clade B) and the six clade C viruses shown in Fig 1 (**Fig 2A, 2B**). EMC and AH13 replicated comparably, as expected, in both A549^DPP4^ and Calu-3 cells. HKU270, an East African clade C strain isolated in Egypt, also replicated fairly comparably to EMC and AH13 in A549^DPP4^ but was attenuated in Calu-3 cells. All other clade C viruses replicated lower than EMC/AH13 in both A549^DPP4^ and Calu-3 cells at all time points, however were only statistically significant at 48hpi. Consistent with the literature [3, 18], this confirms these camel-derived isolates do not replicate as efficiently as wild-type EMC in human lung-derived cell lines. The reduced replication of Mor213 and CAC9690 in Vero-CCL81 cells also suggests these two clade C viruses also contain replication defects that are not necessarily specific to human airway cells.

**Figure 2.**
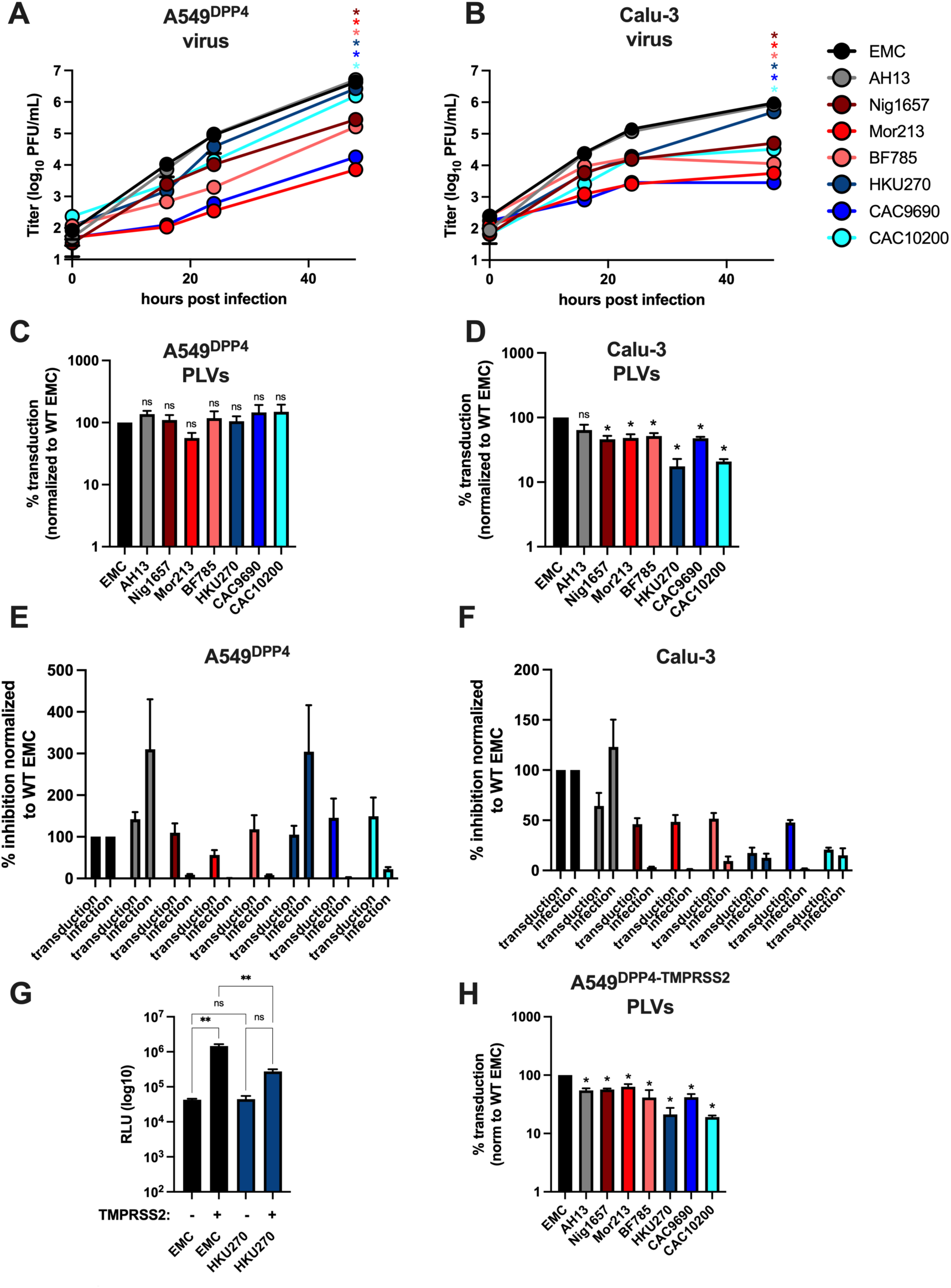
Clade C viruses are attenuated for replication and viral entry in human cell lines. **A,** **B)** A549^DPP4^ (A) or Calu-3 (B) were infected for 16, 24 or 48h (MOI 0.1) and infection quantified by plaque assay. **C, D)** Pseudotyped lentiviral vectors (PLVs) bearing spikes as shown in Fig 1B were used to transduce A549^DPP4^ (C) or Calu-3 (D) and transduction measured by luciferase activity 48h later. Transduction is normalized to that of WT EMC. **E, F)** The relative inhibition, plotted as percentage of WT EMC for infection at 24h or transduction at 48h is plotted for A549^DPP4^ (E) or Calu-3 (F). **G)** A549^DPP4^ were mock or TMPRSS2 transfected 18h prior to transduction with PLVs bearing EMC or HKU270 spikes. **H)** A549^DPP4^ were transfected with TMPRSS2 and transduced with PLVs bearing different spikes 18h later. Transduction was measured by luciferase activity 48h later and normalized to that of WT EMC. All experiments are N=3, *=P<0.05 tested by one-way ANOVA showing statistical significance against WT EMC.

To test whether sequence differences in spike contributed to these attenuated phenotypes, we generated pseudotyped lentiviral vectors (PLVs) bearing the different clade C spike proteins (**Fig 1A**). Transduction of A549^DPP4^ cells with these PLVs demonstrated that, while there were minor differences in transduction efficiency, all clade C spikes were capable of transducing A549^DPP4^ cells to a similar extent (**Fig 2C**). This is consistent with previous reports which found spikes from Morocco, Nigeria, and Burkina Faso transduced 293T-DPP4 cells comparably to, if not better than, wild-type EMC [13]. We next transduced Calu-3 cells and found that Nig1657, Mor213 and BF785 showed a 2-fold decrease in viral entry in these cell lines (**Fig 2D**). Strikingly, however, HKU270 and CAC10200 showed a 5-fold decrease in viral entry relative to EMC. To compare the magnitude of replication versus transduction defect, % inhibition relative to EMC was plotted for each virus or PLV for both A549^DPP4^ and Calu-3 (**Fig 2E, 2F**). This demonstrates that, while all clade C spikes could transduce A549^DPP4^ cells, replication is severely attenuated in this cell line, with the exception of HKU270. In addition to mutations in the spike gene, Nig1657, Mor213 and BF785 also contain deletions of varying extents in ORF4b. Additionally, Nig1657, CAC9690 and CAC10200 all contain a deletion in ORF3. Given the importance of accessory genes in antagonizing innate sensing, it is highly likely that all the clade C viruses in this panel with the exception of HKU270, are attenuated in A549-DPP4 cells due to post-entry effects on viral replication.

However, in Calu-3 cells, the % inhibition of both replication and transduction for HKU270 and CAC10200 was comparable. This suggests that, for HKU270 and CAC10200, the reduced viral replication in Calu-3s can be attributed to defects in viral entry. Given that a major difference in A549 and Calu-3 cells is the presence of TMPRSS2, we hypothesized that the presence of TMPRSS2 in Calu-3 cells and the preferred “early” plasma membrane entry route may explain the difference in viral entry for HKU270 and CAC10200. To test this, we transiently expressed TMPRSS2 in A549^DPP4^ prior to transduction with PLVs bearing the spike of EMC and one of the clade C panel, HKU270 (**Fig 2G**). The presence of TMPRSS2 conferred a 35-fold increase in transduction efficiency for EMC, consistent with previous literature demonstrating EMC spike can utilize the TMPRSS2-mediated pathway for entry [11, 19]. HKU270 PLVs, however, only conferred a non-significant 7-fold increase in transduction, consistent with the reduced transduction of this PLV in Calu-3 cells which endogenously express TMPRSS2. Next, we examined the transduction efficiency of the entire panel of clade C spike bearing PLVs (**Fig 2H**). As expected, the relative transduction efficiency relative to EMC PLVs phenocopied that of Calu-3 cells when TMPRSS2 is present in A549^DPP4^ cells. Altogether, these data suggest that, while there is a post-entry restriction of clade C viruses in human lung cell lines, there may be a difference in viral replication due to differences in the ability to utilize TMPRSS2 for East African clade C viruses.

### Clade C spike proteins are less cleaved at S1/S2 relative to clade A/B spikes

It has previously been reported that cleavage at the S1/S2 of MERS-CoV spike alters entry pathways which may lead to differential viral tropism, with pre-cleavage at the S1/S2 boundary allowing the S2’ site to be cleaved by TMPRSS2 at/near the plasma membrane, and that spikes lacking the furin cleavage site are instead reliant on the endosomal entry pathway for both S1/S2 and S2’ cleavage events [11, 20, 21]. S1/S2 cleavage therefore permits infection of TMPRSS2-expressing cells [11, 22]. As overexpression of TMPRSS2 resulted in reduced transduction by East African clade C PLVs, we hypothesized that the spike aa substitutions present in clade C spikes may alter spike cleavage and therefore TMPRSS2 usage. Thus, we next sought to determine the relative spike cleavage of our viral stocks by western blotting both the cellular lysate and purified viral supernatant after 48hpi in Vero CCL81 (**Fig 3A-3D**). Strikingly, all the clade C spikes appeared to be less well cleaved at the S1/S2 boundary than EMC. Surprisingly, AH13 also appeared to be less cleaved than EMC in this cell-type. We next analyzed the spike cleavage in our PLV producer 293T cellular lysates and purified PLVs (**Fig 3E-H**). In both the cellular lysates and purified PLVs, all spikes were cleaved at the S1/S2 site. While there appeared to be a small difference in S1/S2 cleavage when band intensity was quantified, all spikes were similarly cleaved in the PLV system. Differences in cleavage of coronavirus spike proteins in PLV systems relative to authentic virus, or virus-like particles where E, M and N are also expressed, has previously been reported by other groups [23, 24]. Additionally, SARS-CoV-2 virus-like particles (VLPs) produced in 293T cells compared to Vero cells have been demonstrated to have more variable morphology and differences in spike trimer length [25]. Taken together, we suggest that while useful to interrogate the viral entry step alone, PLV systems may not fully recapitulate authentic differences in spike cleavage in coronaviruses.

**Figure 3.**
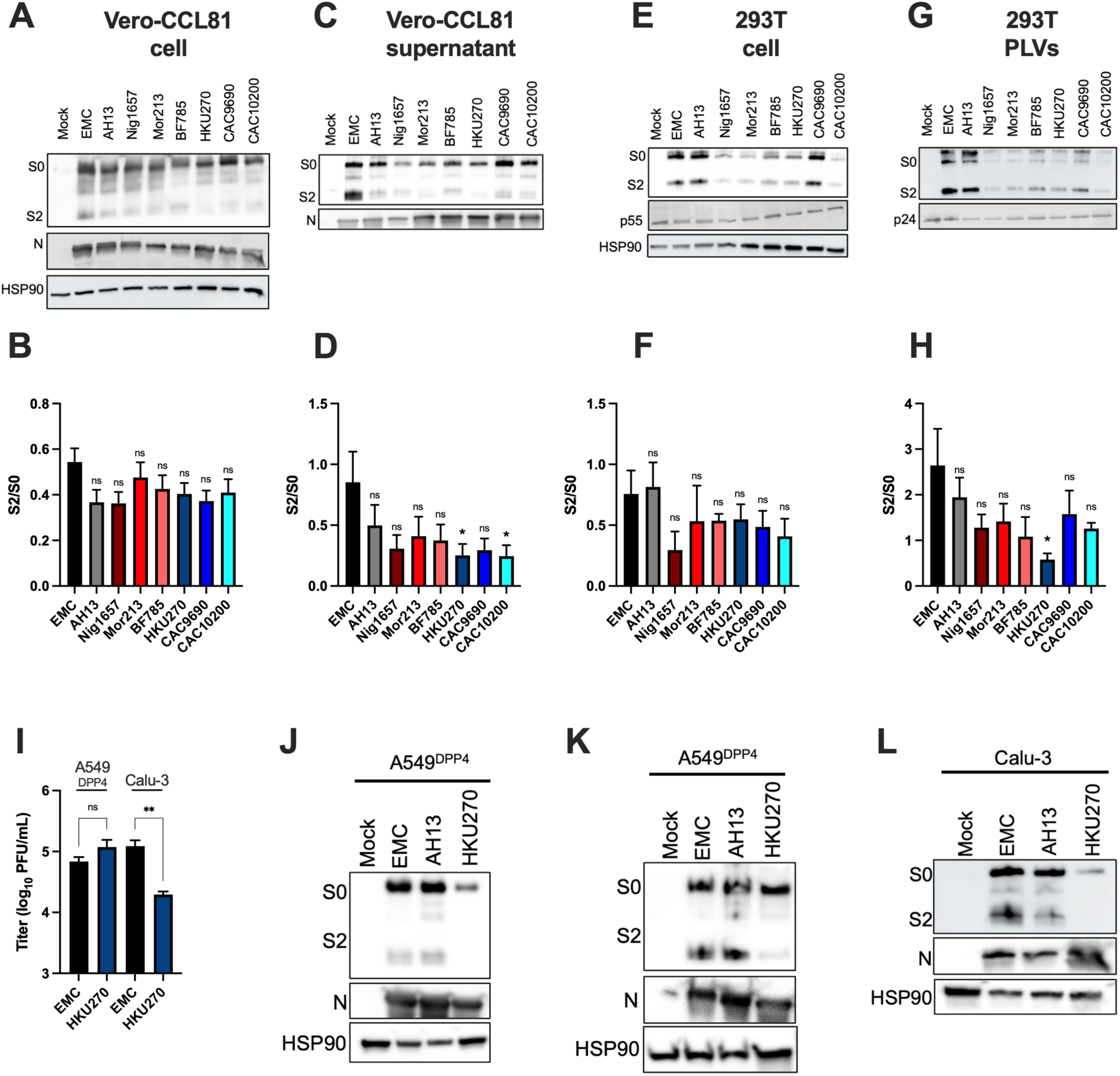
Clade C viral spikes are less cleaved at S1/S2 than WT EMC. A, B, C, **D)** Cellular lysates (A, B) or viral supernatant purified through 20% sucrose (C, D) from Vero-CCL81 and immunoblotted for spike and nucleocapsid. Relative S2 to S0 was quantified in ImageJ. **E, F)** 293T cells transfected to produce PLVs were immunoblotted for spike, p24 and HSP90. Relative S2 to S0 was quantified in ImageJ. **G, H)** PLVs were purified through 20% sucrose and immunoblotted for spike and p24. Relative S2 to S0 was quantified in ImageJ. **I)** EMC and HKU270 were used to infect A549^DPP4^ or Calu-3 cells at MOI 0.1 for 24h and infection quantified by plaque assay. **J)** Cellular lysates of A549^DPP4^ 48hpi were western blotted for spike, Nucleocapsid and HSP90. **K)** Cellular lysates of A549^DPP4^ 48hpi were western blotted for spike and nucleocapsid with 3x as much HKU270 sample loaded onto the gel. **L)** Cellular lysates from infected Calu-3 48hpi were immunoblotted for spike, nucleocapsid and HSP90. All experiments are N=3, *=P<0.05 tested by one-way ANOVA indicating statistical significance against WT EMC.

Due to these differences between spike cleavage during authentic virus infection and PLV generation, we next wanted to analyze spike processing of clade C viruses in A549^DPP4^ and Calu-3 cells. We chose to compare the cleavage of EMC (clade A), AH13 (clade B), and HKU270 (clade C), as HKU270 replicated comparably to EMC and AH13 in A549^DPP4^ but displayed attenuation in Calu-3 at 24hpi (**Fig 3I**). In A549^DPP4^, HKU270 appeared less well cleaved (**Fig 3J**). Interestingly, despite the comparable viral titers in this cell line, we observed less nucleocapsid and spike of HKU270 in the cellular lysates. To confirm that the reduced spike cleavage of HKU270 was not merely an artefact of less detectable spike, more protein of HKU270 was loaded and the spike cleavage again analyzed by western blot (**Fig 3K**). This demonstrated that even with comparable S0 input, HKU270 spike is less cleaved at the S1/S2 boundary in A549^DPP4^ cells. Next, we immunoblotted cellular lysates of Calu-3 cells for spike, and again observed reduced spike expression and cleavage of HKU270 (**Fig 3L**). This suggests that the S1/S2 site is less accessible by furin-like proteases for HKU270 spike in both A549^DPP4^ and Calu-3 cells.

Altogether, these data show clade C spikes are less well cleaved at the S1/S2 junction than clades A and B spikes during native viral infection, despite no sequence differences directly within the S1/S2 cleavage site.

### Clade C viruses are generally reduced for syncytium formation relative to clades A and B viruses

S1/S2 cleavage has previously been shown to be required for efficient cell-cell fusion [26]. Additionally, mutations across both the NTD and S2 of spike have previously been reported to alter syncytia formation of both SARS-CoV-2 variants of concern (VOCs) and MERS-CoV strains [17, 27]. We hypothesized that syncytium formation during infection with these viruses may be altered due to the reduced S1/S2 cleavage of these spike proteins. To test this, we infected A549^DPP4^ cells at MOI 0.1 and stained for nucleocapsid at 16hpi and quantified number of nuclei per syncytium (**Fig 4A, quantified in 4C**). Besides BF785, which formed syncytia comparably to EMC and AH13, all clade C viruses were impaired for syncytium formation. To confirm if the reduced syncytial formation was simply an artefact of differences in replication at 16hpi, we transfected spike alone into A549^DPP4^ cells and again quantified syncytium formation 16h later (**Fig 4B, quantified in 4D**). While the magnitude of difference in syncytium formation was smaller than observed with live viral infection, clade C spikes, with the exception of BF785, still formed syncytia to a reduced extent compared to EMC and AH13. In particular, the HKU270 spike was severely attenuated for syncytium formation. Interestingly, HKU270 replicates comparably to EMC at 16hpi in A549^DPP4^ (**Fig 2A**); however syncytium formation is markedly decreased (**Fig 4**). This suggests the relationship between viral replication and cell to cell spread is not a direct correlation as may be expected.

**Figure 4.**
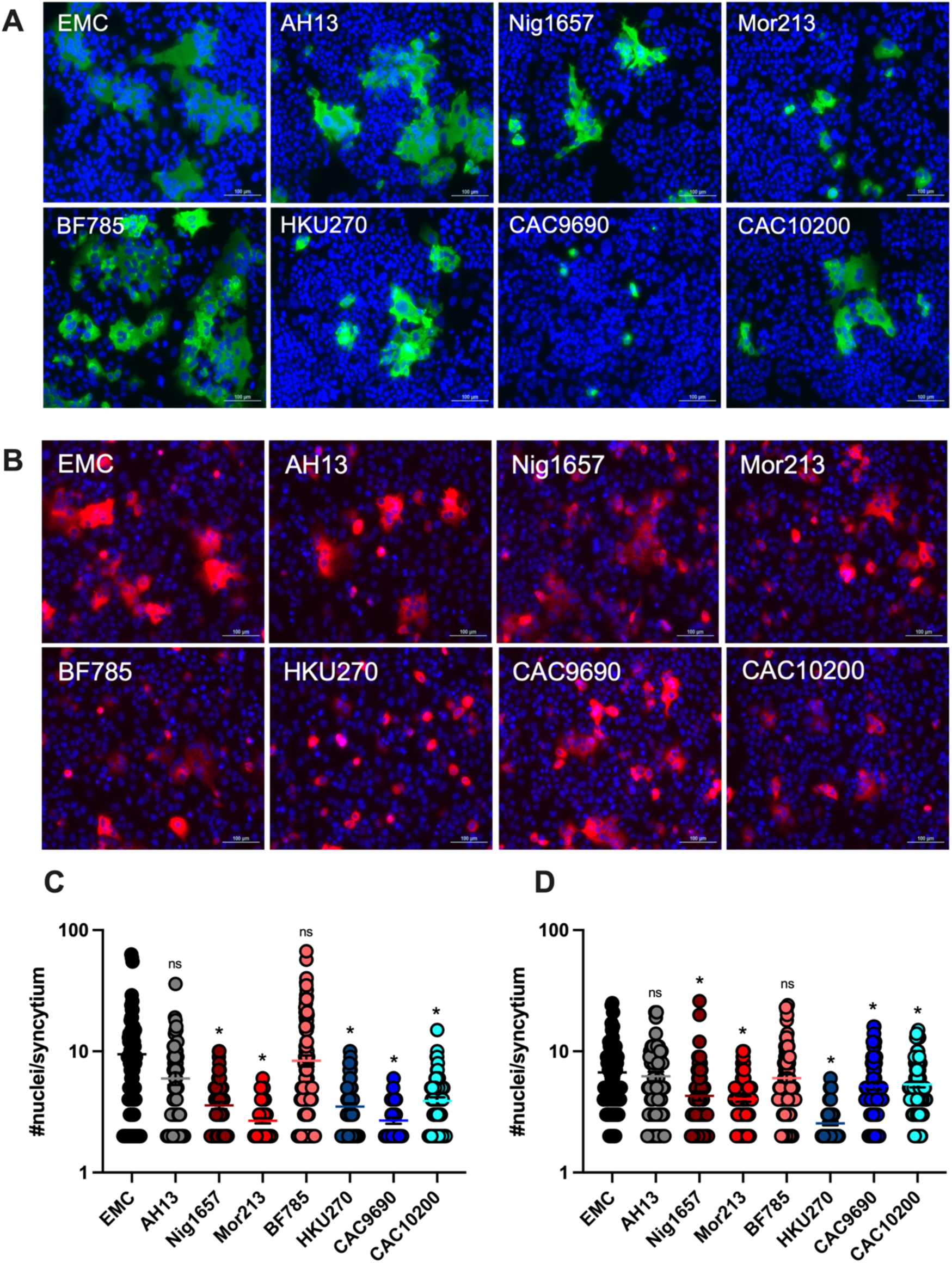
Clade C viruses are varyingly attenuated for cell-cell fusion. **A)** A549^DPP4^ cells were infected at MOI 0.1 for 16h and stained for Nucleocapsid and DAPI. Images were acquired on a Nikon Eclipse Ti 2 and number of nuclei per syncytium counted. **B)** A549^DPP4^ cells were transfected with 500ng of spike plasmid of interest and stained for Nucleocapsid and DAPI 16h later. Images were acquired on a Nikon Eclipse Ti 2 and number of nuclei per syncytium counted. All experiments are N=3 with six images taken per condition per experiment. **C)** Quantification of A. **D)** Quantification of B. *=P<0.05 tested by one-way ANOVA indicating statistical significance against WT EMC.

### Clade C viruses vary in their ability to utilize TMPRSS2 for viral entry

Having established that several clade C viruses appear to be attenuated in a TMPRSS2-dependent manner, we next wanted to investigate if there were differences in the sensitivity of our clade C panel to Nafamostat, a TMPRSS-inhibitor. Calu-3 and A549^DPP4-TMPRSS2^ cells were pre-treated with increasing doses of Nafamostat for 1h prior to infection for 24h with EMC, AH13, and clade C viruses (**Fig 5A, 5B**). EMC and AH13 were very sensitive to Nafamostat pre-treatment in both cell lines, consistent with previous literature on the utilization of TMPRSS-proteases by the EMC strain [11, 19]. West African clade C isolates Nig1657 and Mor213 were also sensitive to Nafamostat pre-treatment. As these viruses also exhibited only a 2-fold reduction in viral entry as PLVs in Calu-3 and A549^DPP4-TMPRSS2^ cells (**Figure 2D**), this suggests that these viruses utilize TMPRSS2 comparably to EMC/AH13 for entry, despite apparently poor S1/S2 cleavage. This could suggest that perhaps minimal S1/S2 cleavage is sufficient for TMPRSS2 usage for these West African viral isolates. BF785 appeared slightly less sensitive than the other West African isolates in Calu-3 cells; however was still inhibited 2-fold by the lowest dose of Nafamostat. In A549^DPP4-TMPRSS2^ cells, BF785 appeared resistant to the lowest dose of Nafamostat, but gained sensitivity comparable to the other West African strains at the highest dose. This could suggest that while BF785 does not utilize TMPRSS2 as well as wild-type MERS-CoV, BF785 can utilize other TMPRSS proteases available in Calu-3 cells that are also inhibited by Nafamostat. The East African clade C isolates, however, which all exhibited the lowest viral entry measured by PLV transduction in Calu-3 and A549^DPP4-TMPRSS2^ cells, were all relatively resistant to Nafamostat until the highest dose in both Calu-3 and A549^DPP4-TMPRSS2^. To confirm that this effect of Nafamostat was only on viral entry and not a post-entry effect of the drug, we tested the sensitivity of clade A, B and C PLVs to Nafamostat in both Calu-3 and A549^DPP4-TMPRSS2^ cells (**Fig 5C, 5D**). Again, East African clade C isolates were more resistant to the effects of TMPRSS-inhibition than EMC, AH13 or West African spikes. Rather than the intermediate phenotype of BF785 found in Calu-3 with live virus, PLVs of BF785 appeared as Nafamostat-resistant as the East African spikes. This suggests that certain spike phenotypes may be difficult to recapitulate with a PLV system. HKU270, CAC9690, and CAC10200 all remained relatively Nafamostat-resistant as PLVs compared to EMC, AH13, Nig1657 and Mor213, confirming these East African spikes are less able to utilize the TMPRSS2-mediated pathway for viral entry.

**Figure 5.**
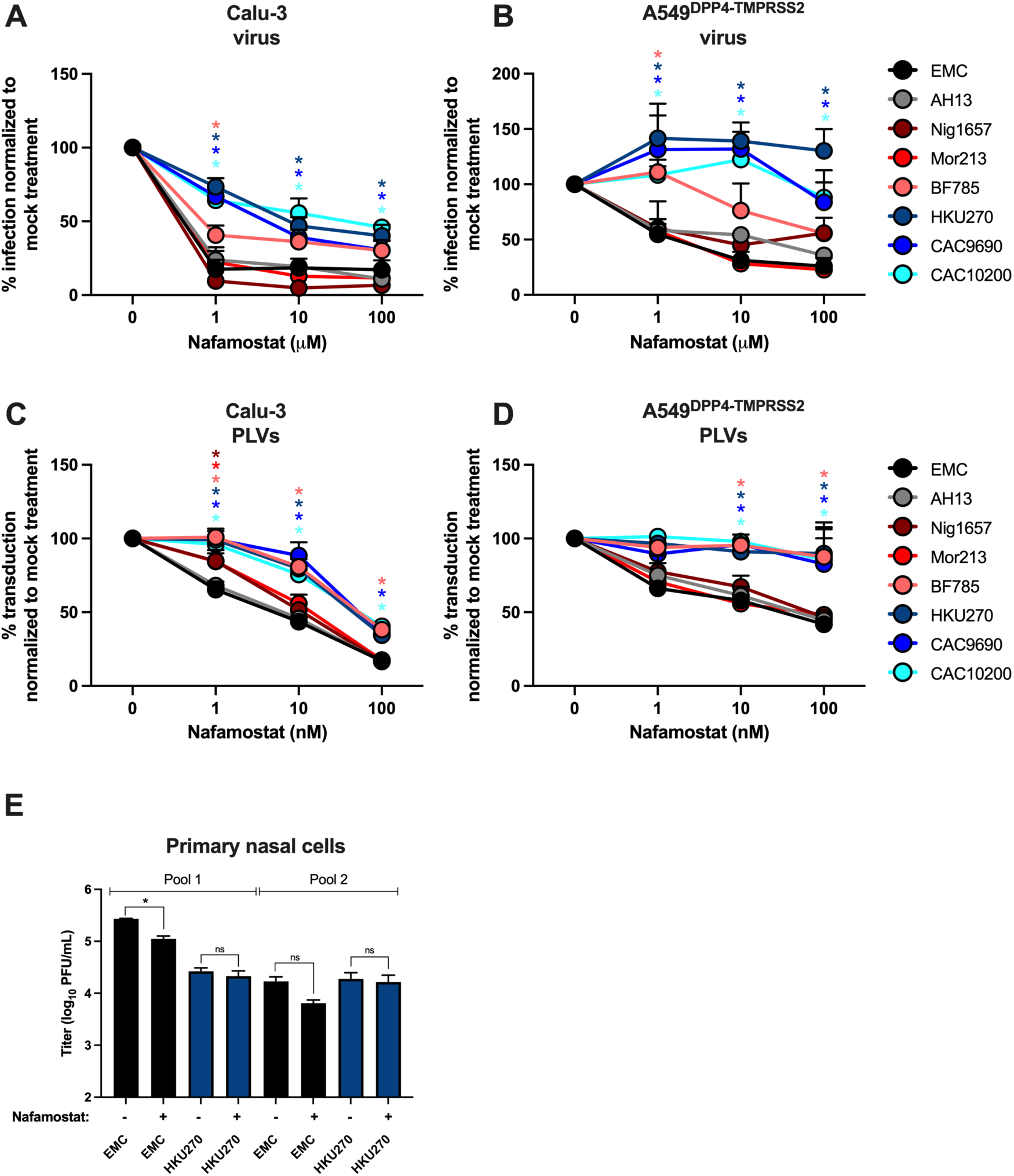
East African clade C viruses are less able to utilize TMPRSS2 for viral entry. A, B) Calu-3 (A) or A549^DPP4^ pre-transfected with TMPRSS2 (B) cells were pre-treated with Nafamostat for 1h prior to infection at MOI 0.1 with viruses indicated. Infection was quantified by plaque assay and infectivity normalized to mock treatment per virus. **C, D)** Calu-3 (C) or A549^DPP4^ pre-transfected with TMPRSS2 (D) were pre-treated with Nafamostat for 1h prior to transduction with PLVs bearing different spikes. Transduction was quantified by luciferase activity 48h later and transduction normalized to mock treatment per spike. Experiments are N=3, *=P<0.05 tested by one-way ANOVA indicating statistical significance against WT EMC. **E)** Primary nasal cells derived from three pooled donors (pool 1 = donors 1627, 1628, 1615, pool 2 = donors 1485, 1578, 1592) were cultured at the air-liquid interface and apically treated with 1μM of Nafamostat for 1h prior to infection with EMC or HKU270 at MOI 1. Two biologically independent infections from 2 independent pools of 3 different donors are shown. *=P<0.05 tested by one-way ANOVA indicating statistical significance between mock treatment and Nafamostat treatment.

While A549 and Calu-3 cells are lung-derived cell lines, we wanted to confirm if differences in TMPRSS2 usage were detectable in primary human nasal cells. Two pooled cultures of primary human nasal cells, each derived from three different donors, were cultured at the air-liquid interface and pre-treated with Nafamostat prior to infection with EMC and HKU270 and infectivity measured by plaque assay 48hpi (**Fig 5E**). Interestingly, we observed differences in the titers of EMC in the mock-treated condition between the two different pools of donors, with pool 1 producing higher titer EMC than pool 2. This may be due to donor-dependent differences in receptor or protease expression, as we observed previously [28]. Despite the differences in titer of the mock-treated condition, EMC was inhibited 3-fold by Nafamostat. HKU270 was, however, entirely unaffected by the addition of Nafamostat, confirming that HKU270 is less able to utilize TMPRSS proteases during infection of primary human airway cultures.

### Both the NTD and SD2 region of HKU270 confer its reduced TMPRSS2 usage

Next, we wanted to determine the mechanism of the reduced TMPRSS2-usage of the East African strains. As shown in figure 3, all clade C strains appeared less well cleaved at the S1/S2 site than EMC. To test if an East African clade C spike could be made to utilize the TMPRSS2-mediated pathway by pre-cleavage at the S1/S2 boundary, we pre-treated EMC and HKU270 with trypsin prior to infection of Calu-3 cells (**Fig 6A**). Trypsin pre-treatment resulted in a modest boost of 2-fold to EMC infectivity, and a 5-fold increase in infectivity of HKU270. Importantly, pre-treatment of both viruses with trypsin prior to infection of Calu-3 cells pre-treated with Nafamostat rendered the previously resistant HKU270 sensitive to Nafamostat (**Fig 6B**). This confirms that HKU270 can utilize the TMPRSS2-mediated pathway if sufficient S1/S2 cleavage occurs. While some of the West African clade C strains were also poorly cleaved at the S1/S2 boundary and yet still capable of utilizing TMPRSS2, this suggests that for HKU270 cleavage at the S1/S2 site is a pre-requisite to TMPRSS2-mediated entry.

**Figure 6.**
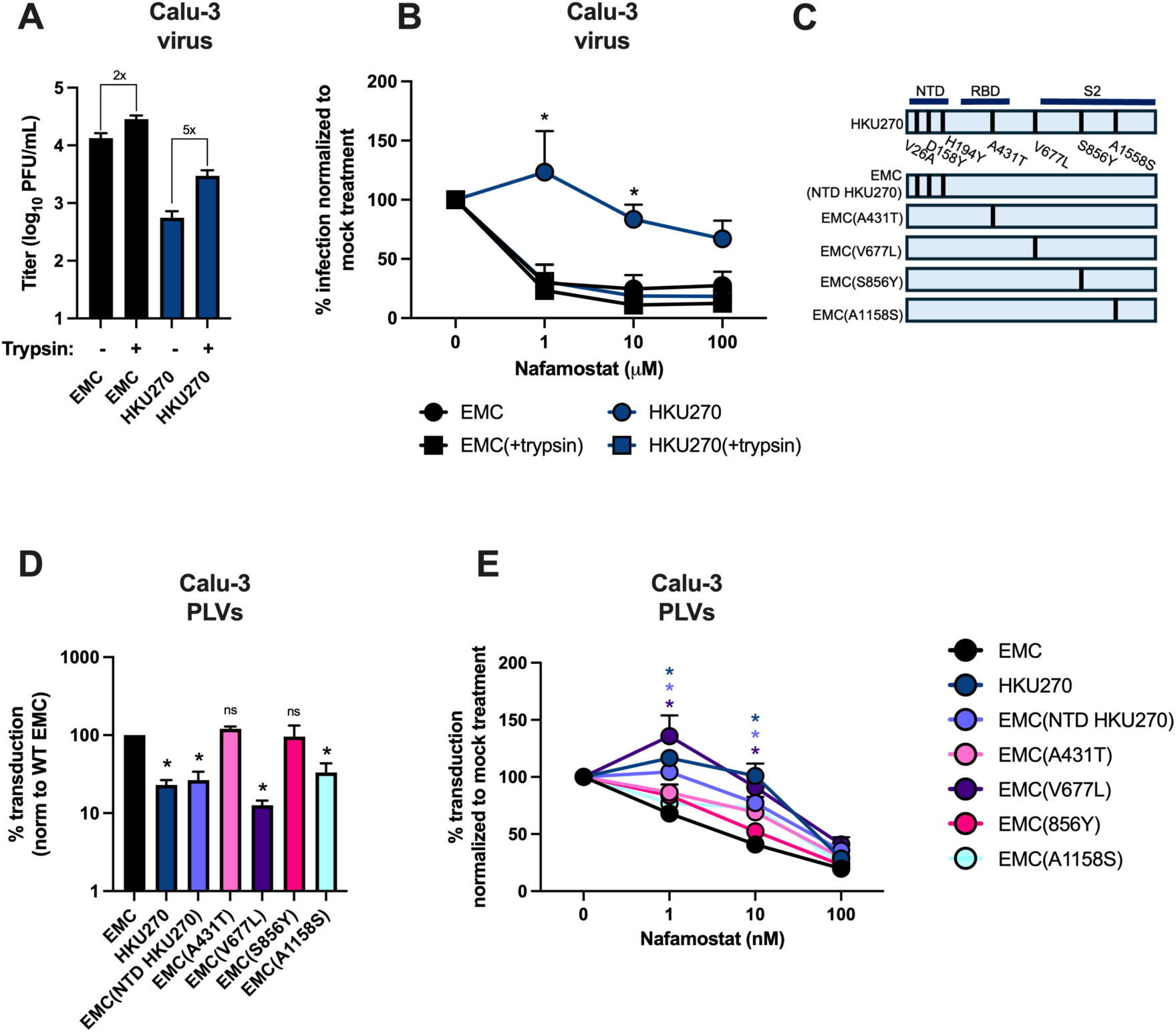
Amino acid substitutions in the NTD and SD2 region of spike govern the reduced TMPRSS2 usage of HKU270. **A)** EMC and HKU270 serum-free stocks were pre-treated with trypsin prior to infection of Calu-3 cells (MOI 0.1) and infectivity quantified by plaque assay 24hpi. **B)** EMC and HKU270 were pre-treated with trypsin as in A and used to infect Calu-3 cells (MOI 0.1) which had been pre-treated with Nafamostat for 1h. Infection was measured by plaque assay 24h later. **C)** Schematic of spike mutants generated into EMC background. **D)** Calu-3 cells were transduced with PLVs bearing different spikes and transduction measured by luciferase activity 48h later. Transduction is normalized to that of WT EMC. **E)** Calu-3 cells were pre-treated with Nafamostat for 1hr prior to transduction with PLVs. Transduction is normalized to mock treatment per PLV. All experiments are N=3, *=P<0.05 tested by one-way ANOVA indicating statistical significance against WT EMC.

Next, we sought to determine the aa substitutions which determine the reduced TMPRSS2 usage of HKU270. HKU270 contains 7 aa substitutions in its spike protein, three in the NTD, one in the RBD, one in the SD2, and two in the S2 domain (**Fig 6C**). Interestingly, the NTD mutations V26A, D158Y and H194Y are identical to that of the two other East African isolates in this study (**Fig 1A**). Next, we generated a panel of spike mutants bearing the V26A/D158Y/H194Y (NTD), A431T (RBD), V677L (SD2), S856Y (S2) or A1158S (S2) substitutions found in HKU270 spike into the background of wild-type EMC spike (**Fig 6C**). Transduction of Calu-3 cells with this panel of mutants demonstrated that both the NTD and V677L substitutions reduced the transduction of EMC spike comparably to that of HKU270 (**Fig 6D**). The A431T substitution did not reduce viral entry at all and transduced comparably to EMC. The A1158S mutation, which is found in all other clade C spikes, reduced viral entry mildly, which may contribute to the mild reduction of viral entry of the West African clade C spikes. Next, we used these PLVs to transduce Calu-3 cells pre-treated with Nafamostat (**Fig 6E**). Again, the NTD and V677L substitutions appeared to confer HKU270-like properties to the EMC spike, with both substitutions separately resulting in reduced Nafamostat sensitivity. This suggests that differing regions of spike can contribute to a reduced ability to utilize the TMPRSS2-mediated pathway during entry.

## Discussion

Here we show that East African clade C isolates HKU270, CAC9690, and CAC10200 are less able to utilize TMPRSS2 for viral entry into cell lines than clade A or B strains, or indeed West African clade C strains. Furthermore, we show that HKU270 is less able to utilize TMPRSS2 in primary nasal cultures compared to the WT EMC isolate. We also identify that aa substitutions in the NTD and SD2 of these East African strain spikes confer reduced TMPRSS2 utilization.

While the West African clade C isolates studied here appear to utilize TMPRSS2 similarly to wild-type EMC and AH13 strains, their spike proteins appear to be poorly cleaved at the S1/S2 boundary. Additionally, all the spike proteins appear more cleaved in 293T cells when generating PLVs than the live virus in Vero-CCL81, A549^DPP4^ or Calu-3, despite differences in protease usage broadly being phenocopied between authentic infection and PLV transductions. Increased spike cleavage in 293Ts is consistent with previous reports on SARS-CoV-2 PLVs versus authentic infection/VLPs [23–25], and is likely explained by differences in protease expression and interactions with other viral structural proteins during viral assembly. Additionally, while betacoronaviruses assemble at the endoplasmic reticulum-Golgi intermediate compartment (ERGIC), PLVs are assembled by HIV gag at the plasma membrane. It is therefore not entirely surprising that these particles exhibit differences in spike cleavage. Despite this more complex relationship between spike cleavage and TMPRSS2 usage, we demonstrate here that the East African clade C virus HKU270 can be sensitized to Nafamostat with pre-treatment with trypsin. Altogether, this suggests that the relationship between S1/S2 cleavage and TMPRSS2 usage is perhaps more complex than previously found. Perhaps subtle differences in S1/S2 cleavage can be sufficient to alter spike conformation and therefore the accessibility of the S2’ cleavage site, which requires further examination and more quantitative proteomics techniques to measure differences in S1/S2 cleavage and spike conformation.

While these viral spikes do not contain any aa substitutions in the S1/S2 site, aa substitutions outside of this region could impact the accessibility of the cleavage site and spike dynamics. It has previously been reported that the constellation of aa substitutions in the NTD of the SARS-CoV-2 early omicron strains resulted in a more “closed” spike that did not adopt the “RBD up” conformation as easily, despite efficient S1/S2 cleavage [15, 17]. Furthermore, it has been reported that the NTD of early omicron spikes partially contributes to the reduced TMPRSS2 usage of omicron, further suggesting that substitutions in the NTD can alter protease usage as demonstrated here [15]. It could be that the sequence differences in the NTD between the West and East African spikes studied here confer different levels of “closed” spike. Of the three substitutions found in the NTD of the HKU270, CAC9690, and CAC10200 spikes, two substitutions (V26A and H194Y) are found in either all or most of the West African isolates. The D158Y substitution, however, is only found in the East African spikes in this panel. Additionally, the D158Y substitution is found in all (41) publicly accessible East African isolates to date but in none (12) of the West African isolates. However, there are less publicly available West African camelid MERS-CoV spikes, and the lack of this mutation found in these sequences so far may be sampling bias. Further sequencing of camel-derived MERS-CoV strains across Africa is needed to further draw conclusions on the geographical distinction of these mutations and protease usage.

The NTD has also been postulated to be important for initial attachment during MERS-CoV entry, in particular in binding to sialic acids [7]. Differences in the NTD sequence of the African MERS-CoV strains may also be due to species-specific differences in sialic acid binding during viral attachment. α2,3-linked sialic acids have been reported to be present in the nasal epithelium of dromedary camels but thought to predominantly be found in the lower airways of humans [8]. Whether differences in sialic acid binding contribute to differential attachment and entry of clade C strains in the human airways remains to be examined and would likely shed light on the different species selection pressures on the NTD of spike.

Interestingly, the V677L substitution in the SD2 region of spike also confers reduced TMPRSS2 usage for HKU270. While the CAC9690 and CAC10200 genomes do not encode this substitution, they encode R626P, which is located just upstream of the SD2 region where V677 is located. The SD2 region of other coronavirus spikes has also previously been implicated in altering protease cleavage at the S1/S2 junction. The D614G aa substitution, which arose early in the SARS-CoV-2 pandemic, was reported to alter furin cleavage efficiency and the up/down ratio of the RBD [29]. The H655Y substitution, found in the omicron SARS-CoV-2 lineage, is also found in the SD2 region and has been reported to increase the preference of omicron on the endosomal route of viral entry and cleavage by cathepsins [14, 15, 17]. Finally, introduction of a Y623H substitution found into SARS-like coronavirus RsSHC014 spike protein increases its entry into human cells by promoting an open spike conformation [30]. While requiring further examination and structural modeling, it is entirely possible that the V677L mutation operates in a similar manner and contributes to reduced TMPRSS2 usage by altering spike conformation. Intriguingly, it has also been reported that the SD2 region responds to the motion of the

NTD and can adopt a different conformation depending on NTD movement, and that small changes in SD2 conformation could lead to large changes in NTD and RBD movement [29]. It may be that the sequence changes in the NTD and SD2 regions of clade C spikes function cooperatively to reduce S1/S2 cleavage and TMPRSS usage.

Somewhat surprisingly, we also find that BF785 is less able to utilize TMPRSS2 than the other West African strains or the WT EMC or AH13. BF785 does not contain an aa substitution in the SD2 region, nor the D158Y substitution found in the East African strains. This suggests there is an alternative mechanism underlying the differential protease usage of BF785 which requires further elucidation. While BF785 appeared moderately sensitive to Nafamostat in Calu-3 cells, BF785 was resistant to Nafamostat in A549^DPP4-TMPRSS2^ cells. This could suggest that while BF785 does not utilize TMPRSS2 it can utilize an alternative serine protease present in Calu-3 cells that is inhibited by Nafamostat [31]. We also show here that clade C spikes, with the exception of BF785, are relatively attenuated for cell-cell fusion compared to clade A and B spikes, even when accounting for differences in viral replication. It is intriguing that HKU270 replicates to comparable titers in A549^DPP4^ cells but does not form as large syncytia; however, BF785 is significantly attenuated for viral replication but forms comparable, if not larger, syncytia than WT EMC. BF785 does contain a large deletion in the ORF4b gene, which encodes NS4b, an antagonist of RNase L and also NFkB signaling [32–35]. The replication of BF785 is therefore likely confounded by a reduced ability to antagonize innate sensing; however, this suggests a somewhat counterintuitive, non-linear relationship between viral output and cell-cell fusion.

It has previously been reported that sequence differences in the NTD of the beta and delta variants of SARS-CoV-2 promote increased cell-cell fusion [17, 36] and that S1/S2 cleavage can be a proxy for prediction of efficiency of syncytium formation. While we find here that BF785 bucks this trend of S1/S2 cleavage, TMPRSS2 usage, and syncytium formation, the three East African clade C viruses studied here perfectly follow this trend of reduced S1/S2 cleavage, reduced TMPRSS2 usage, and reduced syncytium formation. These data also suggest that the BF785 virus exhibits a more cell-associated phenotype than WT EMC, as this virus appears to induce large syncytia, but lower viral titer. This has previously been observed with the mouse hepatitis virus (MHV) strain JHM, where increased receptor-independent cell-cell spread correlates with the increased neurovirulence of this MHV strain [37].

Camels are the amplifying host for MERS-CoV, however the ancestral viruses are believed to have originated in bats, where the viral tropism is suggested to be more enteric than lung-associated [38–45]. The reduced S1/S2 cleavage of these clade C viruses could indicate a preference for replication in the gastrointestinal tract, where cleavage by abundant trypsin-like proteases may facilitate the S1/S2 cleavage event prior to entry. Minimal S1/S2 cleavage during viral production could be advantageous in the gut, to prevent the S2’ cleavage event happening too early before the virus has bound a target cell and spike falling apart. However, it has been shown that WT MERS-CoV replicates in both gut cell lines and primary intestinal epithelial cells, thus the differences in spike cleavage and their relevance to organ tropism require further elucidation [46]. Alternatively, the differences in S1/S2 cleavage observed between clade A/B and clade C spikes could suggest differences in protease availability in the camelid airway relative to humans. Camelid TMPRSS2 has been reported to share 88% homology with human TMPRSS2; however, the prevalence of TMPRSS2 in camel airways has not yet been investigated [47].

An inability to efficiently infect TMPRSS2-expressing cells in the airways could partially explain the attenuation of East African clade C viruses. An increased reliance on endosomal entry could render East African MERS-CoV strains more sensitive to endosomally located antiviral proteins such as interferon-induced transmembrane (IFITM) proteins or nuclear receptor coactivator 7 (NCOA7), which have been shown to inhibit SARS-CoV-2 in an entry route-dependent manner [20, 48]. However, the attenuation of replication of West African clade C isolates appears to be independent of viral entry. Nig1657, Mor213, and BF785 are all attenuated in both A549^DPP4^ and Calu-3 cells, despite robust entry into A549^DPP4^ and only a 2-fold decrease in entry into Calu-3 cells. We suggest that the defects in NS4b of Nig1657, BF785 and Mor213 may impair their antagonism of innate sensing. NS4b has been well characterized as an antagonist of the antiviral protein RNase L, which recognizes and degrades host and viral RNA [32–34]. An inability to disrupt RNase L may explain the reduced replication of West African isolates in A549 and Calu-3 cells observed here. The nuclear localization signal on NS4b has been shown to compete with and prevent nuclear localization of NFkB with potential effects on inflammatory responses [35]. Additionally, Nig1657, CAC9690 and CAC10200 all contain a deletion in NS3, the role of which remains to be examined. Further studies are required to elucidate the entry-independent attenuation of replication of these viruses in human cells.

Altogether, we show here that camel-derived East African isolates of MERS-CoV are less able to utilize the TMPRSS2 protease for viral entry and map the molecular determinants of this differential protease usage. The exact relationship between S1/S2 cleavage and TMPRSS2 usage requires further examination to better understand the reduced S1/S2 cleavage of West African isolates which retain TMPRSS2 usage. However, differences in protease usage could affect the cellular tropism of clade C viruses in the human airways and have consequences for the ability of these viruses to replicate and spread in human tissues.

## Methods

### Cell lines and plasmids

293T (ATCC), Calu-3 (ATCC), and Vero-CCL81 (ATCC) were cultured in Dulbecco’s modified Eagle medium (DMEM) (Gibco) with 10% fetal bovine serum (FBS) and penicillin/streptomycin (Gibco) and incubated at 37°C and 5% CO2. A549^DPP4^, generated as previously described [32] were cultured in RPMI with penicillin/streptomycin at 37°C and 5% CO2.

The EMC WT MERS codon-optimized spike was kindly provided by Nigel Temperton to Stuart Neil who kindly provided it to us. AH13, Nig1657, Mor213, BF785, HKU270, CAC9690, CAC10200 and mutants thereof were generated by Gibson assembly (NEBuilder HiFi Assembly Kit, E5520S, NEB) following the manufacturer’s instructions. CSXW and 8.91 were kindly provided by Stuart Neil. The TMPRSS2 plasmid was kindly provided by Paul Bates.

### Viruses and infection

MERS viruses were kindly provided by Malik Peiris: EMC (GenBank ID: NC_019843.3), AH13 (GenBank ID: KJ650295.1), Nig1657 (GenBank ID: MG923475.1), Mor213 (GenBank ID: MG923469.1), BF785 (GenBank ID: MG923471.1), HKU270 (GenBank ID: KJ477103.2), CAC9690 (GenBank ID: MZ268404.1), CAC10200 (GenBank ID:

MZ268405.1). Viral stocks were generated and titered in Vero-CCL81 cells. Viral stocks were sequenced using Azenta and the following amino acid substitutions found relative to the reference sequences: EMC – orf1a (S1965F, nsp3), spike (T105N), AH13 – spike (A886V, S1251F), M (T8I), Nig1657 – NS5 (P15L), Mor213 – N (K384N), HKU270 – spike (A886V), NS5 (P18L).

For generation of serum-free stocks for trypsin experiments, Vero-CCL81 cells were washed twice with PBS prior to infection with MOI 0.1 of EMC or HKU270 in serum-free DMEM. Cells were again washed twice with PBS 1h post-adsorption and fed with serum-free DMEM. Viruses were harvested 72hpi and titered in Vero-CCL81.

Cells were infected in 2% media, with the exception of trypsin experiments where serum-free stocks were generated and infections carried out using serum-free media and nasal cultures which were infected in serum-free DMEM. Cells were infected for 1h at 37°C and then fed with fresh 2% media for the remainder of the experiment.

For trypsin treatments, serum-free viral stocks were incubated with 3μg/ml trypsin (LS003570, Worthington Biochem) for 12 minutes at 37°C prior to inactivation with soybean trypsin inhibitor (10109886001, Roche) for 5 minutes at room temperature. Viruses were then used to infect cells as per the above and cells fed with 2% DMEM post-adsorption.

### Production of PLVs and transduction

293T cells were transfected with firefly luciferase-expressing vector (CSXW), HIV Gag-Pol (8.91), and spike at a ratio of 3:2:1 μg using 35 μl of PEI-MAX as previously described [20]. Medium was changed 18h later, and vectors were spun at 400 RCF prior to aliquoting and storage at -80°C. 100μl of PLVs were used to transduce cells of interest for 48h and readout measured with the Promega Bright-Glo Luciferase Assay System (E2620, Promega) as per the manufacturer’s instructions on a luminometer (Envision, Perkin Elmer).

### Nafamostat treatment

Cells were pretreated with Nafamostat (30-815-0, Fisher) for 1h at 37°C prior to transduction or infection. Cells were transduced with PLVs for 48h without removing the drug, or media removed and replaced with infection media during viral infections and infection determined 24h later. For Nafamostat treatment of primary nasal cultures, nasal cultures were washed 3x with PBS and treated with 1μM Nafamostat apically in serum-free DMEM for 1h prior to infection.

### Primary nasal ALI cultures

Nasal cells were obtained via cytologic brushing of patients in the Department of Otorhinolaryngology-Head and Neck Surgery, Division of Rhinology at the University of Pennsylvania and the Philadelphia Veteran Affairs Medical Center after obtaining informed consent. The full study protocol, including acquisition and use of nasal cells, was approved by the University of Pennsylvania Institutional Review Board (protocol #800614) and the Philadelphia VA Institutional Review Board (protocol #00781) with off-site waiver approval for BSL-3 work. Patients with history of systemic disease or on immunosuppressive medications are excluded. ALI cultures were grown and differentiated on 0.4 µm pore transwell inserts as previously described [49–51]. Briefly, cytologic brush specimens were dissociated and fibroblast cell population removed, followed by plating onto transwell inserts. Nasal cells were allowed to grow to confluence in the submerged state for approximately 5 days and then apical growth medium removed. Basal differentiation media was replaced three times weekly for 3-4 weeks prior to infection. Basal differentiation medium PneumaCult-ALI basal medium (Stemcell Technologies), was used for all experiments.

### Western blotting

Cell lysates were harvested with RIPA buffer (50mM Tris pH 8, 150mM NaCl, 0.5% deoxycholate, 0.1% SDS, 1% NP40) supplemented with protease inhibitors (Roche, cOmplete mini EDTA-free protease inhibitor) and phosphatase inhibitors (Roche, PhosStop easy pack) and mixed 3:1 with 4x Laemmeli sample buffer (BioRad) prior to boiling at 95°C for 10 minutes. Samples were separated on 4-15% BioRad Mini PROTEAN TGX gels (cat #4561033) prior to transfer onto 0.2uM PVDF membranes. Membranes were blocked in 5% non-fat milk before probing with the following primary antibodies: 1:1000 mouse anti-spike (MA5-29978, Thermo), 1:3000 mouse anti-gag (MA5-44992, Thermo), mouse anti-nucleocapsid (40068-MM10, Sino Biological), anti-HSP90 (Gtx109753, genetex). Proteins were detected using SuperSignal West Femto Substrate (Thermo Fisher, 34095) or SuperSignal West Pico PLUS Chemiluminescent Substrate (Thermo Fisher, 34580) on an ImageQuant Amersham 680. If necessary, blots were stripped using Thermo Scientific Restore Western Blot stripping buffer (catalog #21059) for 30 minutes at room temperature and sequentially probed. For supernatant lysates, supernatant was centrifuged at a relative centrifugal force (RCF) of 20,000 through 20% sucrose for 1h at 4°C prior to harvesting in RIPA buffer.

### Cell-cell fusion assay and immunofluorescence (IF) staining

A549^DPP4^ cells were seeded on glass bottom plates (P24-1.5H-N, Cellvis). For measurement of syncytium formation from live virus infection, cells were infected at MOI 0.1. 1h post-adsorption, cells were fed with 2% media and incubated at 37°C 5% CO2 for 16h. Cells were fixed with 4% paraformaldehyde and removed from the BSL-3. For measurement of syncytium formation by spike-only transfection, cells were transfected with 500ng of each spike using lipofectamine 2000 (Invitrogen). 16h post-infection or transfection, cells were fixed in 4% paraformaldehyde at room temperature for 30 minutes. Cells were washed with PBS and permeabilized with 0.1% Triton X-100 in PBS for 15 minutes at room temperature, washed with PBS and subsequently blocked with 3% BSA in PBS for 30 minutes at room temperature. Cells were incubated with anti-Nucleocapsid (40068-MM10, Sino Biological) or anti-spike (MA5-29978, Thermo) in 3% BSA for 2h at room temperature with rocking. Cells were washed 3x with PBS and incubated with secondary Alexa Fluor antibodies in 3% BSA for 1h at room temperature with rocking. Cells were again washed 3x and DAPI in PBS added for 5 minutes at room temperature. DAPI was washed off with PBS and images acquired on a Nikon Ti2 Eclipse at 20x magnification. Six images per condition were acquired and nuclei per syncytium counted.

All data is available upon request.

## Acknowledgements and funding.

We are grateful to Malik Peiris for providing viruses and to Stuart Neil, Nigel Temperton, and Paul Bates for providing plasmids. We thank Andrew Marques for assistance with analysis of viral sequencing. We thank all members of the Weiss group for their feedback and discussion.

This work was funded by NIH grants R01 AI140442 to S.R.W, R01 AI169537 to S.R.W and N.A.C, and VA grant BX005432 to N.A.C.. H.W. is supported in part by the Brody Family Medical Trust Fund Fellowship in “Incurable Diseases” of The Philadelphia Foundation. H.S. is supported in part by The Martin and Pamela Winter Infectious Disease Fellowship Program, University of Pennsylvania Institute for Infectious & Zoonotic Diseases.

## Author Contributions

Research was designed by H.W., R.A.P.M and S.R.W. Experiments were performed by H.W. Data was analyzed by H.W. Viruses were grown by H.W., H.S., and R.A.P.M. Primary nasal cultures were generated by L.H.T. N.A.C. provided reagents and funding. H.W. wrote the manuscript. All authors edited the manuscript and provided comments.

